# Human Neural Organoid Microphysiological Systems Show the Building Blocks Necessary for Basic Learning and Memory

**DOI:** 10.1101/2024.09.17.613333

**Authors:** Dowlette-Mary Alam El Din, Leah Moenkemoeller, Alon Loeffler, Forough Habibollahi, Jack Schenkman, Amitav Mitra, Tjitse van der Molen, Lixuan Ding, Jason Laird, Maren Schenke, Erik C. Johnson, Brett J. Kagan, Thomas Hartung, Lena Smirnova

**Affiliations:** Center for Alternatives to Animal Testing (CAAT), Johns Hopkins University, Baltimore, MD; Department of Environmental Health and Engineering, Johns Hopkins University, Baltimore MD; Cortical Labs Pty Ltd; Melbourne, Australia; Department of Electrical and Computer Engineering, Princeton University, Princeton NJ; Department of Physics and Astronomy, Johns Hopkins University, Baltimore MD; Neuroscience Research Institute, University of California Santa Barbara, Santa Barbara, CA; Department of Molecular, Cellular and Developmental Biology, University of California Santa Barbara, Santa Barbara, CA; Research and Exploratory Development Department, Johns Hopkins University Applied Physics Laboratory, Laurel, MD, United States; CAAT-Europe, University of Konstanz, Konstanz, Germany; Department of Biochemistry and Pharmacology, University of Melbourne, Parkville, VIC 3010, Australia; Doerenkamp-Zbinden Chair for Evidence-based Toxicology, Department of Environmental Health and Engineering, Johns Hopkins University, Baltimore MD

## Abstract

Brain Microphysiological Systems including neural organoids derived from human induced pluripotent stem cells offer a unique lens to study the intricate workings of the human brain. This paper investigates the foundational elements of learning and memory in neural organoids, also known as Organoid Intelligence by quantifying immediate early gene expression, synaptic plasticity, neuronal network dynamics, and criticality to demonstrate the utility of these organoids in basic science research. Neural organoids showed synapse formation, glutamatergic and GABAergic receptor expression, immediate early gene expression basally and evoked, functional connectivity, criticality, and synaptic plasticity in response to theta-burst stimulation. In addition, pharmacological interventions on GABAergic and glutamatergic receptors, and input specific theta-burst stimulation further shed light on the capacity of neural organoids to mirror synaptic modulation and short-term potentiation, demonstrating their potential as tools for studying neurophysiological and neurological processes and informing therapeutic strategies for diseases.

**Graphical Abstract:** 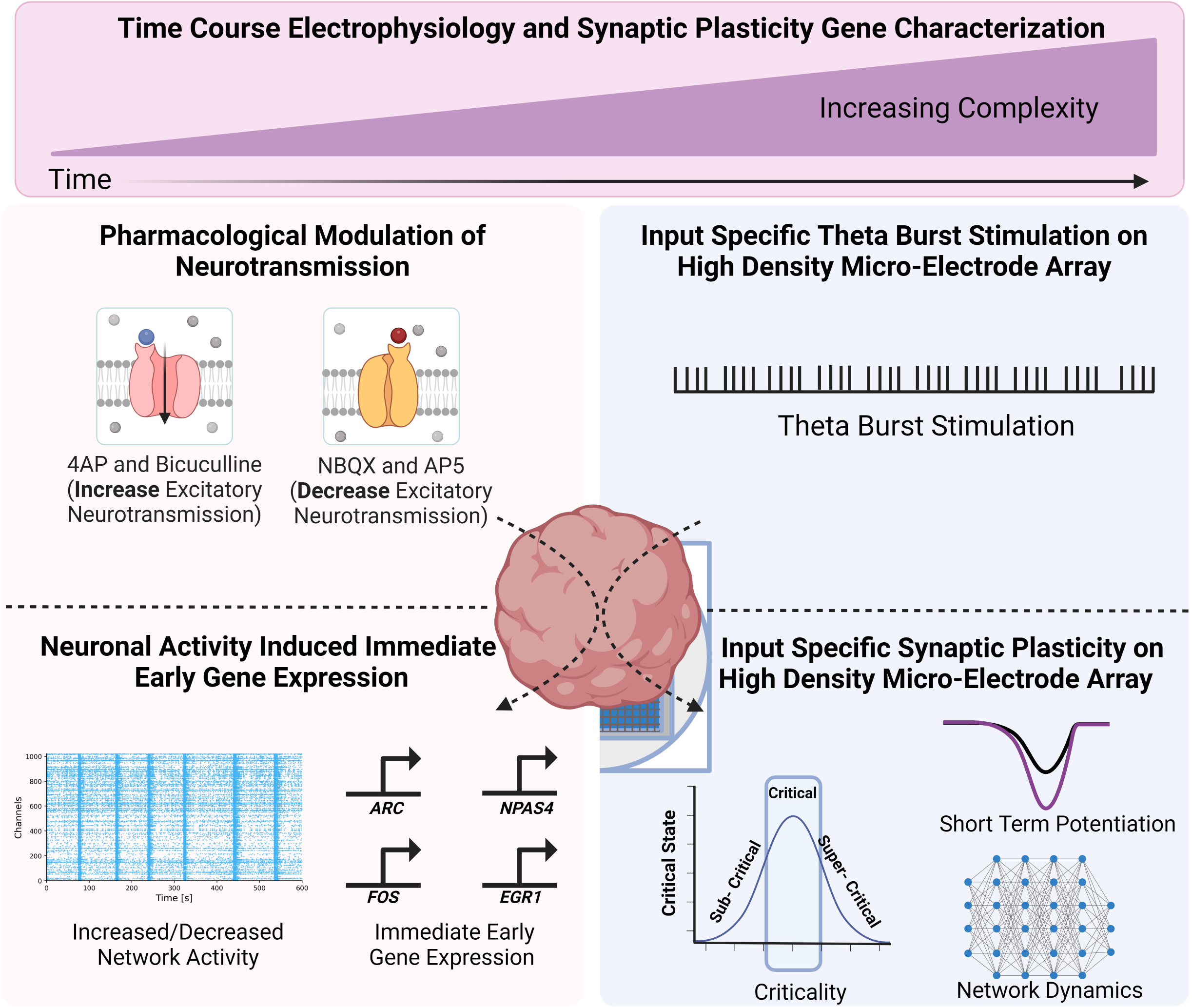

Overview of the main components of the experiments conducted. Figure created using BioRender.com.

## Introduction

Neural organoids; which can be grouped under the umbrella term of Brain Microphysiological Systems (bMPS), are derived from human induced pluripotent stem cells (hiPSCs) and offer a powerful tool for studying brain development, disease modeling, drug discovery, and personalized medicine^1,2^. They can recapitulate key features of the human brain, including cellular diversity^3–6^, connectivity^7^, and functionality^3,4,8^ and can capture specific donor genotypes^9^. The model used for this study is comprised of all the neural cell types found in the human brain including GABAergic, glutaminergic, cholinergic, dopaminergic neurons; neural progenitor cells (NPCs), astrocytes, and myelinating oligodendrocytes^10,11^. Functional analysis of another organoid model has shown that their oscillations are similar to the human preterm neonatal EEG features^12^. Other organoid models have been shown to harbor neuronal assemblies with ample size and functional connectivity, enabling them to collaboratively trigger field potentials^7^. Recently, neural organoids have been proposed as a model of cognition, potentially capable of modelling learning and memory (OI – organoid intelligence)^13^. Towards this goal, organoids were used for reservoir computing, demonstrating spatial information processing and network plasticity as a form of unsupervised learning^14^, but the extent to which bMPSs model the mechanisms of learning and memory have not been fully explored. To develop a reliable learning-in-a-dish model, we first need to understand the molecular and cellular machinery of learning, neuronal network activity and function, and synaptic plasticity in neural organoids, which is extensively characterized here.

One critical aspect of brain functionality that is important for learning and memory, is synaptic plasticity^15–18^. Short- and long-term potentiation (S(L)TP) are activity-dependent forms of synaptic plasticity associated with short- and long-term learning and memory processes^15,19,20^. LTP occurs at the cellular level and involves modifications in synaptic transmission to enhance signal conduction^15,21,22^. Both LTP and STP are NMDA receptor-dependent forms of synaptic plasticity^23^. Synaptic plasticity relies on the rapid activation of immediate early genes (IEGs) in response to stimuli and plays a key role in mediating the transcription and translation of other genes involved in the formation and maintenance of memories^24^. Altered synaptic plasticity leads to abnormal neural network activity, impairing cognitive function and behavior and has been linked to various neurological and psychiatric disorders^18,25^.

Criticality is another important aspect of neuronal activity that has been shown to optimize the ability of neuronal networks to encode and process information^26^. At the critical state, neuronal activity exhibits scale-free dynamics, allowing for efficient information processing and integration across different brain regions^27^. In addition, research has shown that criticality is important for learning and memory in the brain^28^. Research in monolayers of cortical cultures suggests that criticality may be a fundamental property that arises in dynamic systems receiving structured information, making it a valuable metric to assess in more complex cultures^27^. Despite this perspective, aspects of criticality in neural organoids are less explored^29,30^.

The capacity of hiPSC-derived neural organoids to demonstrate features of synaptic plasticity is still not fully understood. One study has shown that organoids respond to external electrical signals and maintain elevated neuronal activity short term^31^. Another study has shown that assembloids exhibit STP/LTP using patch clamp methods^32^. Additionally, Zafeiriou et al., showed that neuronal organoids exhibit both short- and long-changes in network dynamics^33^. While these are great first steps, more work needs to be done to characterize and better understand synaptic plasticity in neural organoid models.

Here, we focus on analyzing the foundational elements of learning and memory in neural organoids through the characterization of spontaneous and evoked neural network dynamics, input-specific synaptic plasticity, functional connectivity, IEG expression, and criticality. We show that our model has spontaneous and evoked highly interconnected neural networks that change over time, show expression and activation of IEGs, and demonstrate critical dynamics.

## Results

Neural organoids were differentiated from iPSCs for up to 14 weeks and characterized throughout development (Fig. 1A). Gene expression of synaptic plasticity markers was quantified from week 0 to week 12. Calcium signaling development was analyzed from week 2 to week 14. Finally, electrical activity was characterized by high-density microelectrode arrays (HD-MEAs) over two independent time periods, from weeks 6-to-9 and 10-to-13. In addition, pharmacological modulation of neurotransmission was performed at week 8 and 13. Lastly, theta burst stimulation (TBS) was implemented at week 13 to induce synaptic plasticity. To analyze both spontaneous and evoked electrical activity from the HD-MEA data, functional connectivity and criticality analysis were performed. A schematic overview of the neurocomputational investigations is shown in Fig. 1B. In addition, an example of how evoked activity from pharmacological or electrical stimuli can modulate synaptic transmission to induce synaptic plasticity is shown in Fig. 1C.

**Figure 1.**
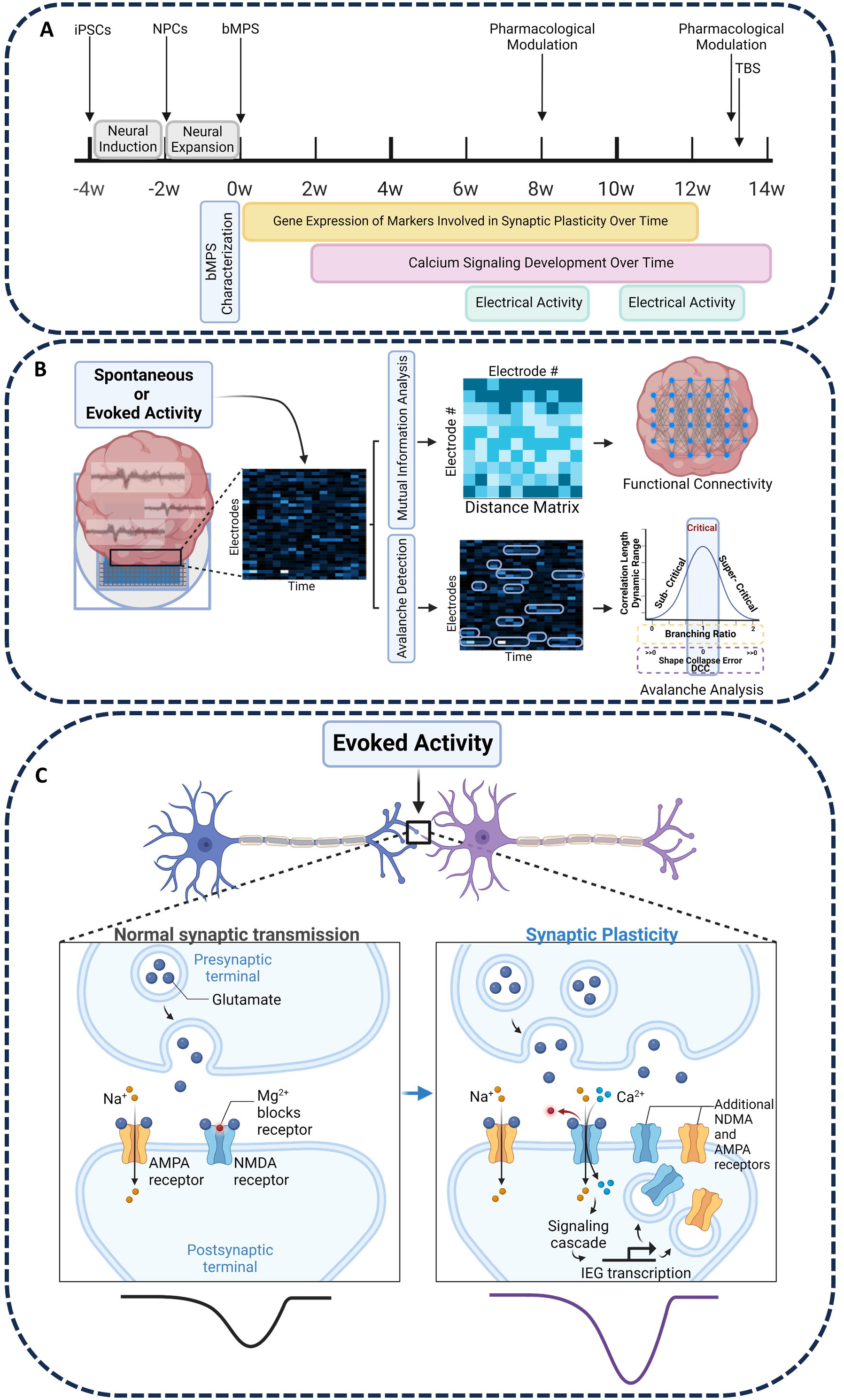
Schematic overview of the experimental approach. A) Experimental timeline. B) Overview of avalanche and network connectivity analysis for time series electrophysiology data from organoids plated on HD-MEAs. C) Schematic representation of synaptic transmission modulation by pharmacological and electrical stimuli to induce synaptic plasticity. Figure created using BioRender.com.

### Neural Organoids Develop Proper Synapse Formation and Express Receptors Necessary for Synaptic Transmission

Neural organoids were differentiated following our in house established protocol^11^. The expression of markers for astrocytes (*GFAP)*, oligodendrocyte (*MBP*, *OLIG2)* and mature neurons (*MAP2)* increased in the first 8 weeks of maturation and then plateaued in the following weeks, indicating that the differentiation predominantly occurs rapidly until week 8 and then reaches a more stable, mature cell composition from week 8 to 12 (Fig. S1). Hence, two time points were selected for the experiments in this paper.

To determine which brain region best matches the neural organoids, RNA-seq data from neural organoids differentiated for 8 weeks were deconvoluted using the BrainSpan developmental transcriptome^34^. Deconvolution using multiple methods converged on the temporal neocortex and ventrolateral prefrontal cortex (Fig. S1).

Expression of the postsynaptic marker *HOMER1* and presynaptic marker Synaptophysin (SYP) was detected in both week 8 and 12 organoids (Fig. 2A). Gephyrin-positive signal was close to background with few positive cells at week 8 and increased at week 12 (Fig. 2B). This indicates that there are more developed inhibitory synapses at the later stage of differentiation. Gene expression of *GABRA1* which encodes the inhibitory GABA_A_ receptor followed the same pattern (Fig. 2C). Gene expression of postsynaptic marker *HOMER1* was steady over time (Fig. 2C).

**Figure 2.**
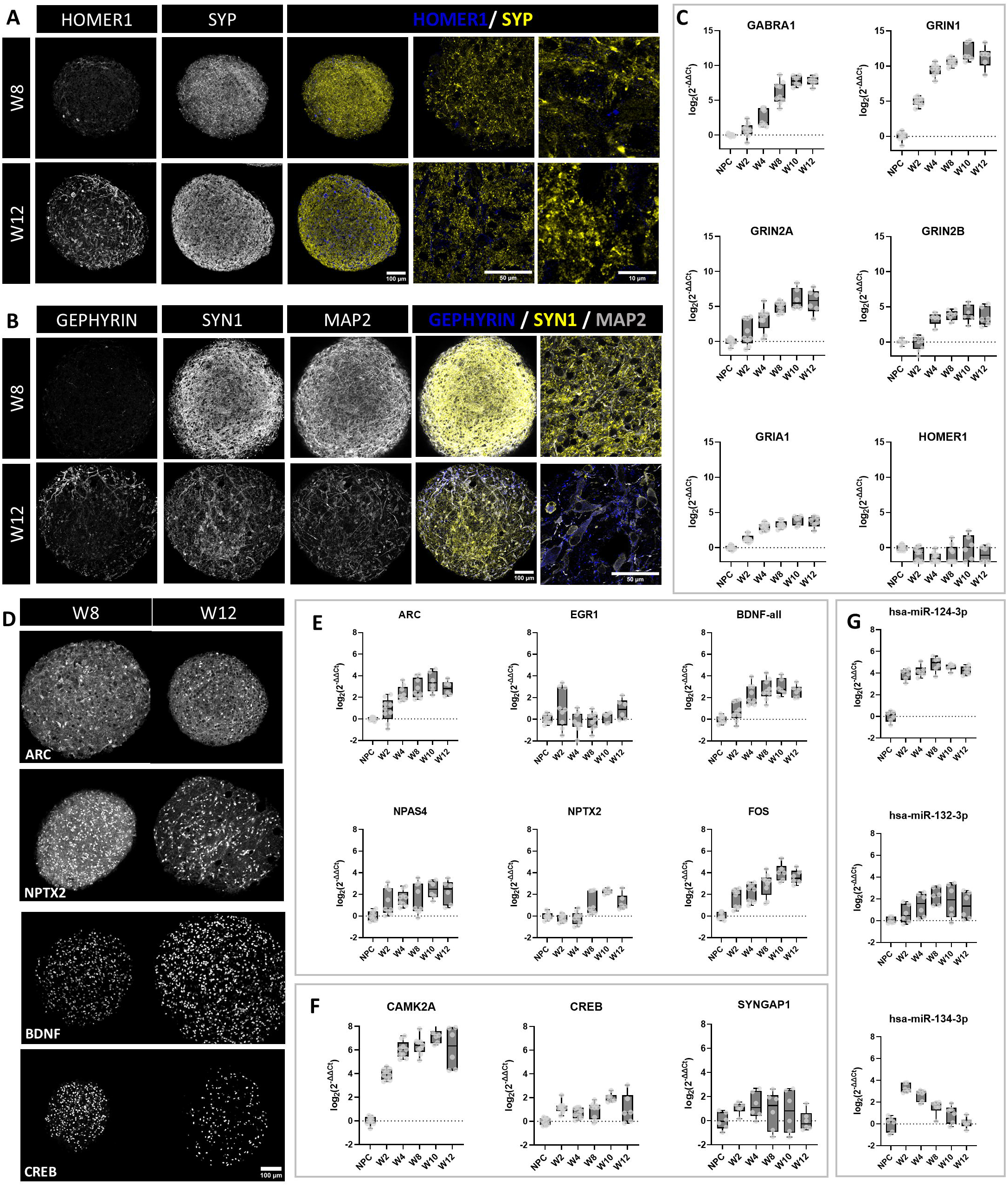
Expression of glutamatergic and GABAergic receptor and synaptic plasticity related genes in neural organoids over course of differentiation. A) Representative immunocytochemistry images of organoids showing postsynaptic marker (HOMER1) and presynaptic marker (SYP) in 8- and 12-week cultures. In composite images, HOMER1 is shown in blue, and SYP is shown in yellow. Scale bars are 100 µm, 50 µm, and 10 µm respectively. B) Presence of inhibitory post-synaptic marker (Gephyrin), pre-synaptic marker (SYN1) and dendrites (MAP2) in 8- and 12-week organoids. In composite images, Gephyrin is shown in blue, SYN1 in yellow, and MAP2 in grey. Scale bars are 100 µm and 50 respectively. For panels A-B) All images were taken at 20x, 100x, and 100x + 4x zoom and processed with ImageJ for visualization. C) Gene expression of Gamma-Aminobutyric Acid Type A Receptor Subunit Alpha1 (*GABRA1*), Glutamate Ionotropic Receptor NMDA Type Subunit 1 (*GRIN1*), Glutamate [NMDA] Receptor Subunit Epsilon-1 (*GRIN2A*), and Glutamate [NMDA] Receptor Subunit Epsilon-2 (*GRIN2B*), Glutamate Ionotropic Receptor AMPA Type Subunit 1 (*GRIA1*), homer scaffold protein 1 (*HOMER1*) in organoids over the course of differentiation. D) Representative immunocytochemistry images of weeks 8 and 12 organoids stained for Neuronal Pentraxin 2 (NPTX2), Activity -Regulated Cytoskeleton-associated protein (ARC), cAMP response element-binding protein (CREB), and Brain-Derived Neurotrophic Factor (BDNF). Scale bar is 100 µm. E) Gene expression over the course of differentiation of immediate early genes (IEG) *ARC*, *BDNF*, Neuronal PAS Domain Protein 4 (*NPAS4*), *NPTX2*, and Fos proto-oncogene AP-1 transcription factor subunit (*FOS*), and Early Growth Response 1 (*EGR1*). F) Gene expression of Synaptic Plasticity Related Genes *CREB*, calcium/calmodulin-dependent protein kinase II A (*CAMK2A*), Synaptic Ras GTPase-activating protein 1 (*SYNGAP1*). G) Gene expression of Synaptic Plasticity Related miRNA. For all gene expression plots, data is shown as box and whiskers plot (extending from the 25th to 75th percentiles) and represented as log_2_(FC) normalized to NPCs from 2-3 independent experiments with 3 technical replicates each. In all qPCR experiments, ACTB was used as a reference gene.

Both AMPA and NMDA receptors play an important role in synaptic plasticity including STP/LTP^23,35^, therefore showing expression of these receptors was imperative for this study to give insight into the mechanisms of learning and memory in neural organoids. The increase in gene expression over time was the greatest for *GRIN1*, which plateaued around week 8 to week 12 (Fig. 2C). *GRIN2A* and *GRIN2B* both increased over time with a higher increase of *GRIN2A* expression than *GRIN2B*, suggesting increasing maturity^36^(Fig. 2C). *GRIA1* expression also increased over time and plateaued after week 8 (Fig. 2C). Thus, plateau in expression of these subunits suggests the organoids reached a mature state between week 8 to 12.

### Dynamic Expression of Immediate Early Genes Associated with Synaptic Plasticity and Cognitive Functions Over Time

IEG are crucial for cognitive functions as they act directly at the synapse and mediate the cellular processes that are essential for learning and memory consolidation^24^. To demonstrate that the neural organoids have the necessary cellular components for cognitive processes, we quantified IEG expression during the course of differentiation (Fig. 2D and E). Gene expression of *ARC*, *BDNF*, *NPAS4*, *NPTX2*, *FOS* was significantly increased over time, while *EGR1* was already expressed in NPCs and remained at levels close to those in NPCs. Expression of upstream regulators of IEGs (*CREB* and *CAMK2A*) also increased over time with the largest increase in expression in *CAMK2A* (Fig. 2F). In addition, *SYNGAP1*, which plays a key role in regulating synaptic function and plasticity^37^ was stably expressed throughout the course of differentiation, starting from NPCs. The levels of IEGs proteins (NPTX2, ARC, and BDNF) and upstream IEG transcription factor *CREB* were comparable between week 8 and 12 confirming the plateau observed in RT-qPCR data (Fig. 2D).

Finally, we assessed the expression of microRNAs known to be involved in synaptic plasticity (Fig. 2G)^38^. A strong increase in expression of *mir-124* was observed. *mir-132* and *mir-134* had opposite expression patterns: *mir-132* was increased over time while expression of *mir-134* was first strongly induced from NPC to 2 weeks of differentiation and was downregulated thereafter (Fig. 2G).

### Evidence of Spontaneous Electrical Activity and Highly Interconnected Neuronal Networks in Neural Organoids

Electrophysiology over the course of organoid development was characterized using calcium imaging and HD-MEAs. In addition to the expression of molecular machinery involved in synaptic plasticity, neural organoids showed highly patterned spontaneous electrical activity (Fig. 3 and Fig. 4). Calcium transients were measured using Fluo-4 biweekly from week 2 to 14. Whole organoid change in fluorescence over resting fluorescence intensity (delta F/F) was quantified and compared across age groups (Fig. 3A). From these plots, the average rise time, peak amplitude, firing rate, decay time, burst duration, number of peaks, and percentage of active organoids per time point were calculated (Fig. 3B). Bursts were identified as peaks in calcium transients. Burst firing rate was calculated as the number of burst peaks per second.

**Figure 3.**
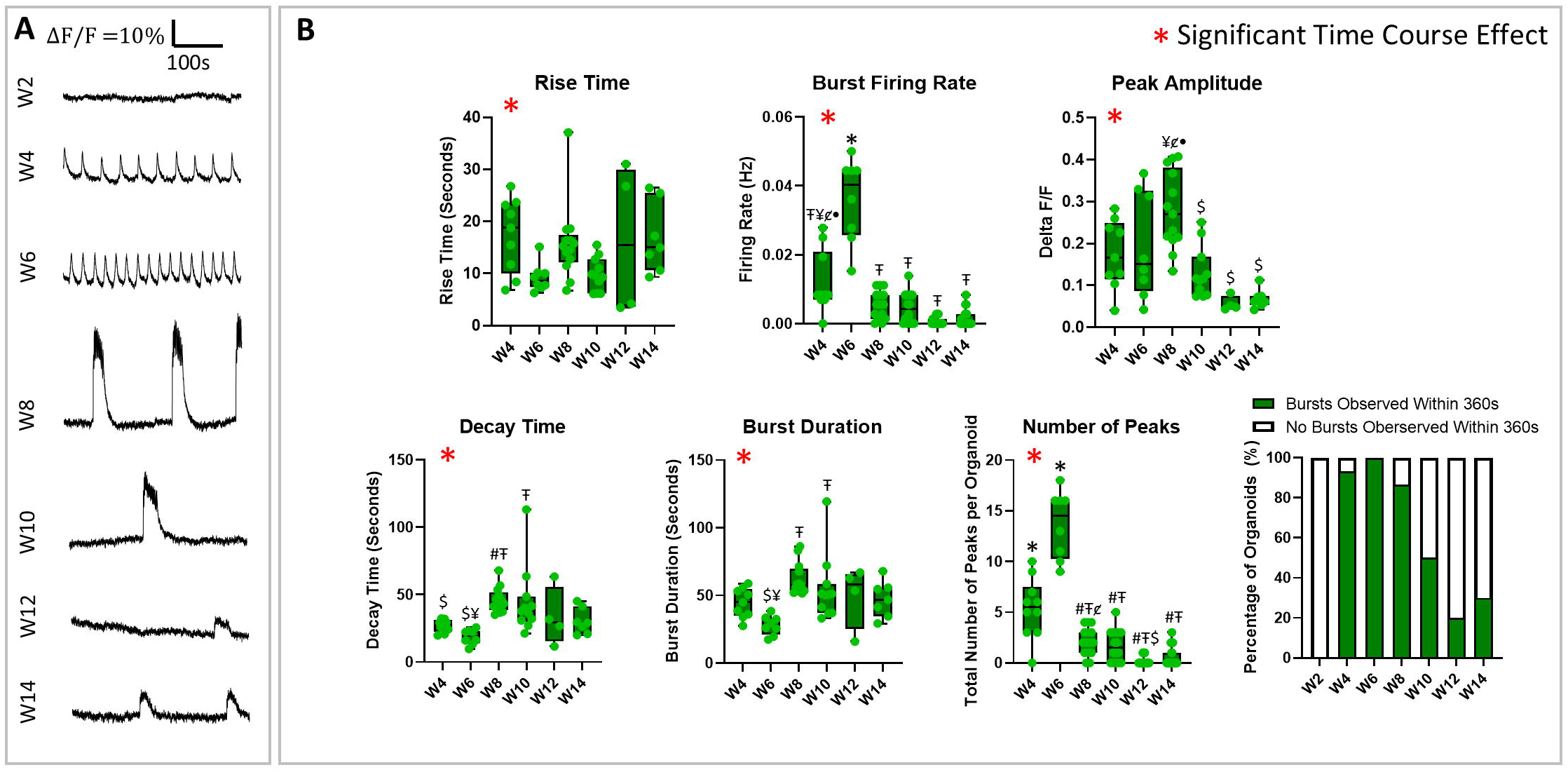
Neural organoid calcium oscillatory dynamics across different time points to show maturation of spontaneous network bursting. A) Representative changes in fluorescence over resting fluorescence (ΔF/F) graphs across 360 seconds for each time point. B) Average rise time, peak amplitude, firing rate, decay time, burst duration, number of peaks, and percentage of active organoids shown across different time points. At least 8 individual organoids across at least 3 independent experiments were imaged and quantified for each time point. Statistics was performed using One-way ANOVA and a Tukey post-hoc test. Changes over time were significant for rise time (p < 0.05), burst firing rate (p<0.0001), peak amplitude (p<0.0001), decay time (p < 0.01), burst duration (p < 0.001), and total number of peaks per organoid (p<0.0001). Pairwise comparisons are shown on the figure: # = Significant difference from week 4, Ŧ = Significant difference from week 6, $ = Significant difference from week 8, ¥ = Significant difference from week 10, lil = Significant difference from week 12, • = Significant difference from week 14, * = Significant difference from all weeks. For exact p values see supplementary tables 4-8. See also Figure S2 for single neuron calcium imaging analysis.

**Figure 4.**
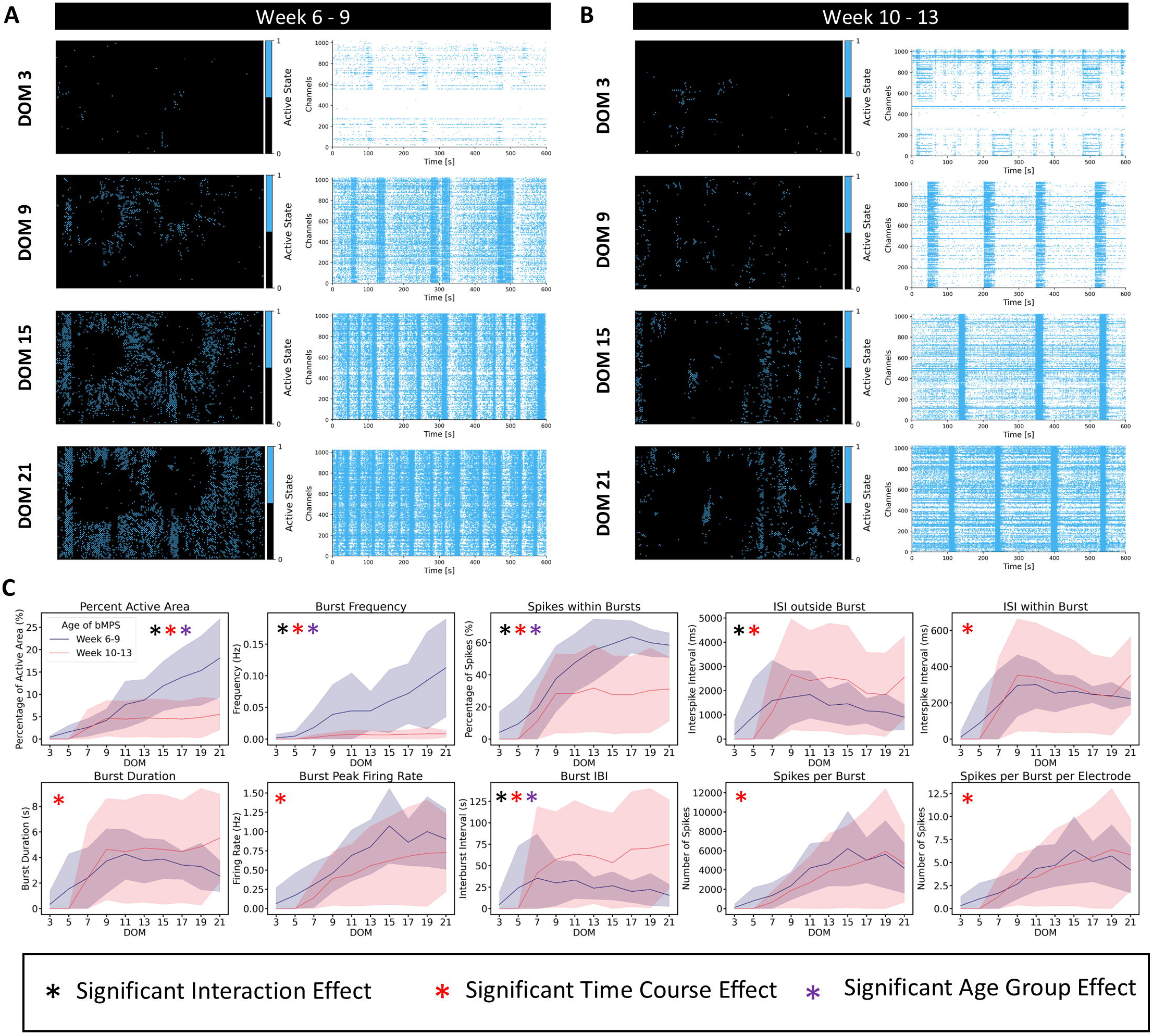
Changes in spontaneous electrical activity in neural organoids throughout development. Representative raster plots and Active Area plots from HD-MEAs recording showing spontaneous electrical activity over time during (A) weeks 6-9 and (B) weeks 10-13 of differentiation. DOM: Days on MEAs. C) Network dynamic metrics from both organoid age groups seeded on HD-MEA over time. Data shown represents mean and standard deviation plotted from 2 independent experiments with 5 to 6 HD-MEA wells per group per experiment with 2-5 organoids per well. Statistics was performed using a mixed-effects model with matching and a Tukey post-hoc test. P<0.05 was considered significant. For exact p values from pairwise comparisons see supplementary documents. ISI: Interspike Interval. IBI: Interburst Interval.

Calcium imaging showed that 2-week-old organoids did not exhibit spontaneous oscillatory calcium dynamics. The first signs of oscillatory calcium activity were detected first at week 4, with high-frequency oscillations at weeks 4 and 6, as shown by higher burst firing rates and number of peaks (Fig. 3, Vid. S1, and Fig. S2). At week 8, oscillation patterns shifted to lower frequency with fewer calcium peaks, lower burst firing rates, higher amplitudes, longer burst durations, and larger decay times (Fig. 3, Vid. S2, and Fig. S2). The plateau shape of the oscillations at week 8 indicated multiple neuronal action potentials contributing to the calcium oscillation (Fig. 3A). The decrease in the number of peaks from week 6 to week 8 suggested more synchronous calcium transients, indicating a more densely connected mature network. From weeks 10 to 14, burst duration, decay time, and number of peaks did not change significantly, but amplitude and percentage of active organoids decreased, suggesting a plateau in differentiation around week 8.

In addition to whole organoid analysis, delta F/F was quantified in single neurons for weeks 4-10 (Fig. S2). Maximum intensity z-projections of time course videos showed that neuronal networks at weeks 4 and 6 were less connected compared to weeks 8 and 10 (Fig. S2). Neurons spiked at higher frequencies and with less synchronization (Fig. S2A and Fig. S2B). By weeks 8 and 10, larger burst amplitudes contributed by multiple action potentials across different neurons, which were spiking simultaneously (Fig. S2C and Fig. S2D). At week 10, the propagation of an action potential across connected neurons was observed by the slightly delayed peak burst amplitude of region of interest (ROI) 1 compared to ROI 2 and 3 (Fig. S2D).

To measure network activity over time, HD-MEAs were used to obtain high spatial and temporal resolution of organoids’ electrical activity across two different time periods (weeks 6-to-9 and 10-to-13) (Fig 4). Representative raster plots indicated differences in spontaneous electrical activity in organoids depending on their age (Fig. 4A-B). The 6-to-9-week organoids have a significantly higher burst frequency, number of spikes within burst, and percent active area than the later time point group (Fig. 4C). They also had significantly shorter inter-burst intervals compared to the more mature group, consistent with the calcium imaging data in Figure 3.

To further assess the organoids’ functionality, neuronal connectivity and criticality were quantified from the same HD-MEA time course data (Fig. 5 and Fig. 6). In both age groups, changes in functional connections between electrodes can be observed over time on the HD-MEA (Fig. 5A). More dense connections and active electrodes were observed in the 10-to-13-week group compared to the 6-to-9-week group as denoted by the thickness of the black lines and red electrodes respectively in the connectivity graphs (Fig. 5A). However, while both groups showed significant increases in the number of nodes over time, the 10-to-13-week group had a significantly lower number of nodes overall in their functional connectivity matrices compared to the 6-to-9-week group (Fig. 5B). To quantify the shift in the strength of the edges over time, an edge weight distribution was calculated by measuring the fraction of total possible edges that is realized (Fig. 5C). Interestingly, most edges are activated across all samples over time (Fig. 5C). The 10-to-13-week group shows no significant changes over time while the 6-to-9-week group shows a temporary significant decrease in strength of edges at DOM 7 (Fig. 5C). Finally, the organoid’s modularity was significantly different across age groups and significantly decreased in both age groups over time, indicating that the networks started with multiple communities but then became more of a single community over time (Fig. 5D and Fig. S3). The decrease in modularity may also be due to an increased number of nodes. Despite the similarity in modularity, the 10-to-13-week group maintained a significantly higher modularity over time, indicating that it maintained more communities or network connections (Fig. 5D).

**Figure 5.**
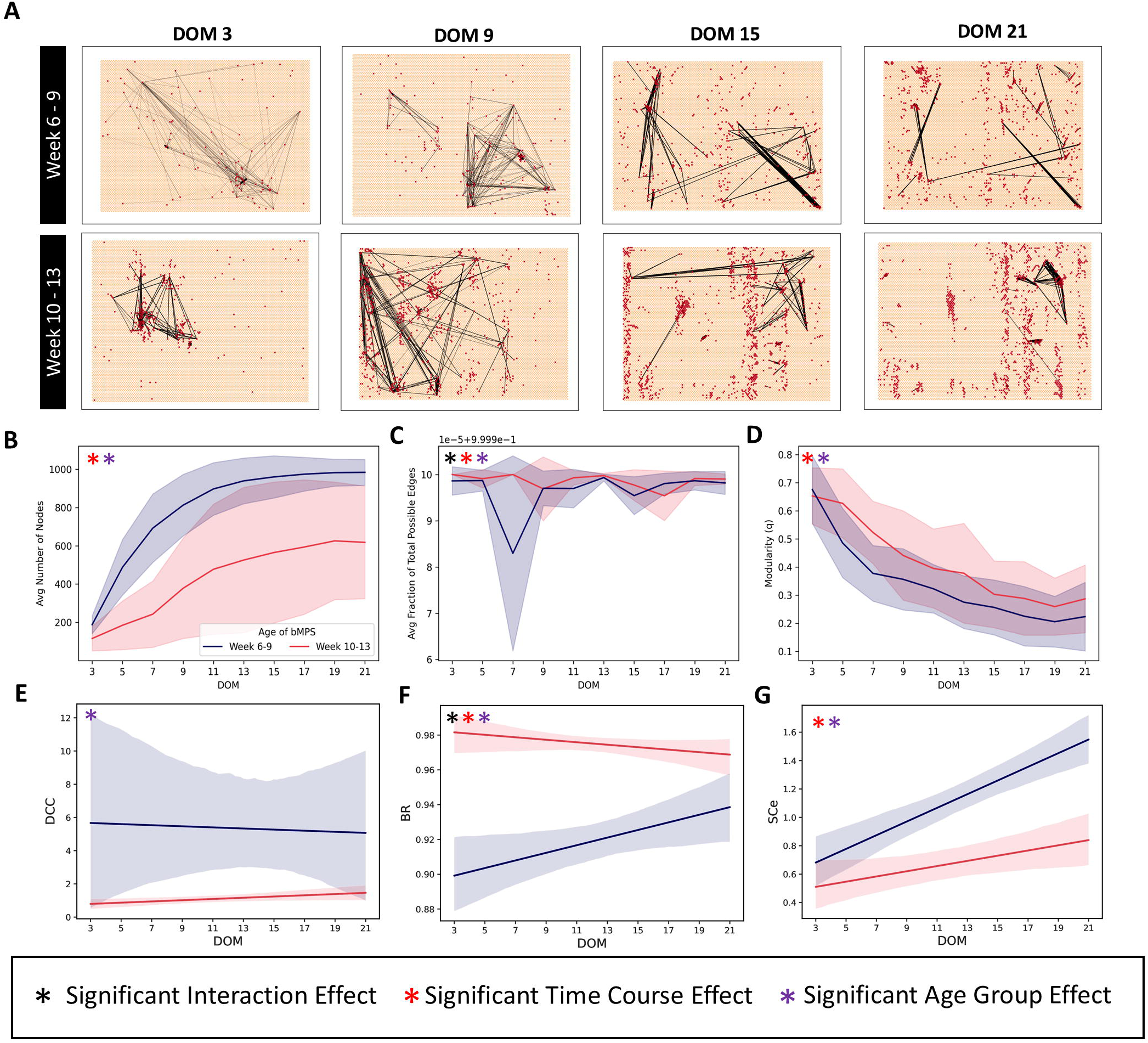
Neural organoids show highly interconnected neuronal networks and criticality throughout development. A) Representative plots of functional connectivity at DOM 3, 9, 15, and 21 for the week 6-9 and week 10-13 old organoids. For clarity of visualization, only the 200 connections (edges) with the highest mutual information are shown. Each red dot represents an electrode, and the lines indicate the connections between electrodes. The thickness of the line indicates the weight of connectivity. See also Figure S4 for an expanded version of network connectivity across all days on the MEA. B) Average number of nodes; C) Average Fraction of Total Possible Edges; D) Average modularity over time in week 6-9 and week 10-13 organoids. E) Number of criticality coefficient (DCC) F) Branching ratio (BR) G) Shape collapse error (SCe) over time in 6-9 week and 10-13 week old organoids. Panels B-D show mean and standard deviation. Panels D–G show regression lines with a 95% confidence interval. Data plotted is from 2 independent experiments with 5-6 HD-MEA wells per group per experiment. Statistics were performed using a 2-way ANOVA and a Tukey post-hoc test. P<0.05 was considered significant. For exact p values from pairwise comparisons see supplementary documents.

**Figure 6.**
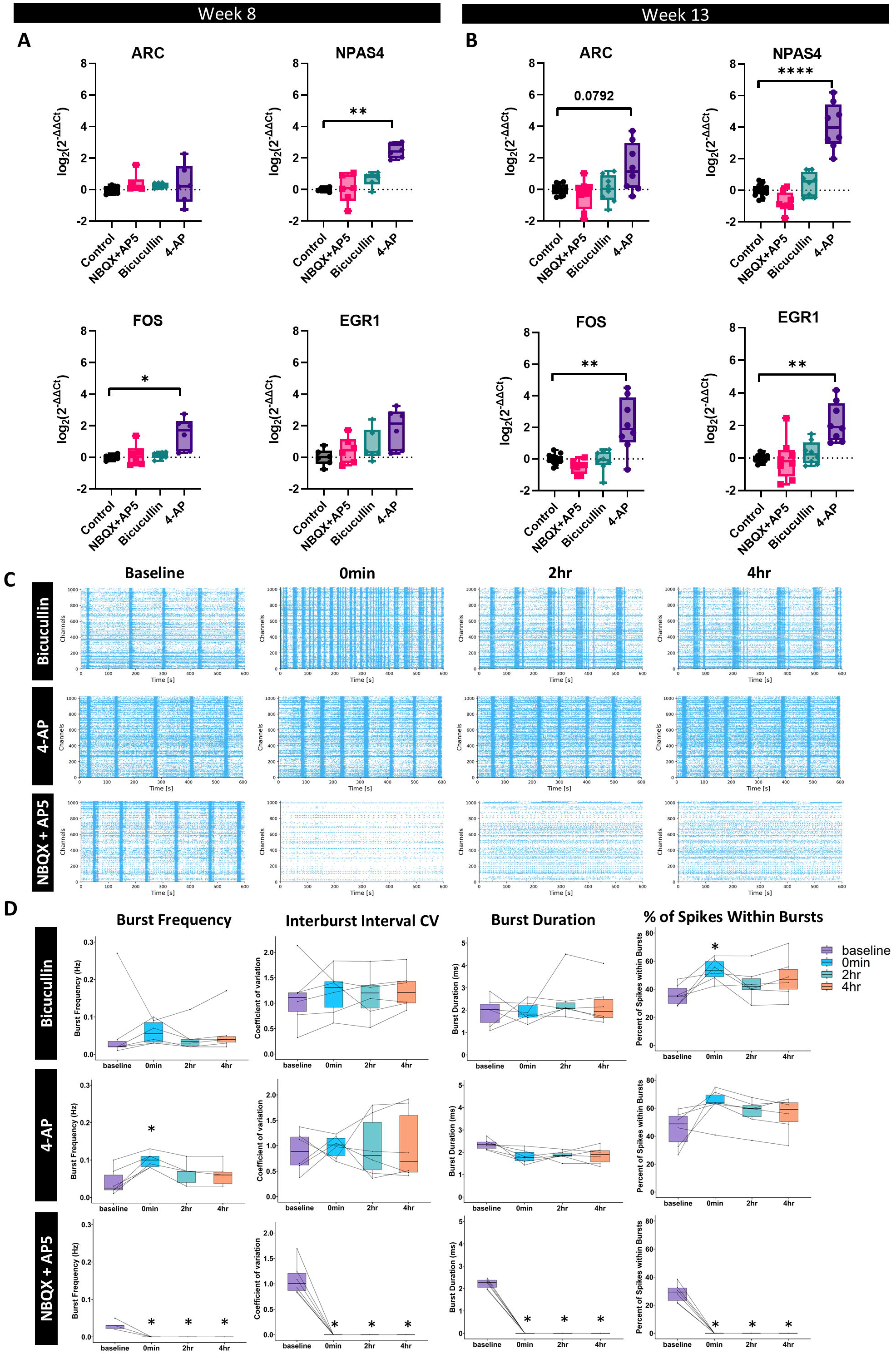
Pharmacological characterization of synaptic transmission changes of neuronal spiking and bursting activity and Immediate Early Gene expression. Expression of *ARC, NPAS4, FOS*, and *EGR1* after 2 hours of exposure to 20 µM AP5+20 µM NBQX, 10 µM bicuculline and 100 µM 4-AP in (A) 8-week and (B) 13-week old organoids, represented as box and whisker plots (25th to 75th percentiles) and as log_2_(Fold Change) normalized to negative control (organoids with no chemical treatment = 2h Control). *ACTB* was used as a reference gene. Data represents 3 independent experiments with 2 technical replicates each for 8-weeks and 4-5 independent experiments with 2 technical replicates each for 13-week time point. Statistics were calculated based on the technical replicate average from each independent experiment, with one-way ANOVA and post-hoc Dunnett’s tests *p < 0.05, ***p < 0.001, ****p < 0.0001 C). Representative Raster Plots from MEA recordings in week 13 old organoids (from 6 wells per condition) before and after treatment with bicuculline, 4-AP, and NBQX+AP5. D). Burst Frequency, Interburst interval coefficient of variation, burst duration, and percentage of spikes within bursts plotted for Bicuculline, 4-AP, and NBXQ+AP5 treated wells before, 0 mins, 2 hours, and 4 hours after exposure. Data represents 3 independent experiments with 2 HD-MEA wells per experiment per chemical (n=6). Statistical significance was calculated with repeated measures ANOVA with post-hoc Dunnett tests. p<0.05 was considered significant. Pairwise comparisons can be seen in the supplementary tables 9-19 and significant groups are shown in the figure.

Criticality is a state of complex systems such as a brain, which operates at the critical point between organization and randomness, demonstrating how neuronal network may navigate between the two stages of chaos and order^39^. The critical point state is a key for brain functionality, as at this stage it operates at its optimal and most efficient computational capacity and is highly sensitive to external stimuli. Organoids exhibited properties of criticality over the course of differentiation (Fig. 5E-G). The more mature 10-to-13-week group showed a consistently lower and more tightly regulated Deviation from Criticality Coefficient (DCC) value and higher branching ratio (BR, approaching 1) compared to the 6-to-9-week group (Fig. 5E). While the BR in the 10-to-13-week group decreased non significantly over the period of 3 weeks on the HD-MEAs, the 6-to-9-week group gradually increased significantly, demonstrating maturation and pursuit of criticality thus stable state (Fig. 5F). Additionally, the shape collapse error (SCe) for the 10-to-13-week group is significantly lower than that of the 6-to-9-week group, indicating a more accurate scaling of avalanches of varying durations to a universal shape in the 10-to-13-week group (Fig. 5G). This analysis suggests that the 10-to-13-week group is in a more critical state compared to the 6-to-9-week group. However, over time, both the BR and SCe appear to converge for both groups, suggesting that the 6-to-9-week group exhibited increasingly critical dynamics, while the 10-to-13-week group showed diminishing critical dynamics on the MEA over time.

### Pharmacological Characterization of Synaptic Transmission Changes Neuronal Bursting Activity and Immediate Early Gene Expression

To validate reactiveness to network modulations, pharmacological agents were used to cause neuronal depolarization and disrupt excitatory glutamatergic synaptic transmission. Expression of IEG and synaptic plasticity-related genes was measured 2 hours after exposure to pharmacological agents and compared to the corresponding untreated control in two age groups (8 weeks and 13 weeks) (Fig. 6). To disrupt excitatory glutamatergic synaptic transmission, organoids were treated with 2,3-dioxo-6-nitro-7-sulfamoyl-benzo[f]quinoxaline (NBQX), an AMPA receptor antagonist, D-2-amino-5-phosphonovalerate (AP5) a NMDA receptor antagonist. 4-Aminopyridine (4-AP), a voltage-gated potassium (Kv) channel antagonist, and bicuculline, a GABA receptor antagonist, were used to enhance neuronal depolarization and synaptic transmission (Fig. 6).

Bicuculline showed a slight increasing trend in gene expression across both age groups (Fig. 6A), while exposure to 4-AP led to significant changes in *NPAS4* and *FOS* expression at both age groups. Expression of *ERG1* was significantly induced only at week 13. Lastly, *ARC* expression showed an increased trend in expression after 4-AP exposure (Fig 6A). No significant changes in gene expression were seen after exposure to NBQX and AP5 individually or combined (Fig. 6 and Fig S3).

Since IEG were more strongly perturbed at week 13, the effects of these chemicals on electrophysiological activity were assessed in this age group. Organoids were exposed to the pharmacological agents directly on the HD-MEA at day on MEA (DOM) 29. Network recordings were taken before the addition of the chemicals as a baseline. Network activity was then recorded immediately after, 2, and 4 hours after the exposure and recorded parameters were compared to baseline activity (Fig. 6C and D). 4-AP and bicuculline increased network activity while NBQX+AP5 decreased network activity over time (Fig. 6C). More specifically, bicuculline caused an insignificant increasing trend in mean burst frequency and interburst interval coefficient of variation (CV) over time, a significant increase in percent of spikes within bursts 0 minutes after and increasing trend in percent of spikes within bursts 2 and 4 hours after exposure. In addition, bicuculline caused no significant changes or trends in burst duration over time. 4-AP exposure caused a significant increase in mean burst frequency and increasing trend in mean percent of spikes within bursts after 0 minutes after. In addition, the percentage of spikes within bursts maintains the increasing trend within 2 and 4 hours after exposure. 4-AP also caused a decreasing trend in burst duration that is maintained over time. Finally, 4-AP caused no significant changes or trends in interburst interval CV over time. Additionally, NBQX/AP5 exposure completely abolished network bursting activity (Fig. 6C and 6D, Fig. S3). Overall, NBQX+AP5 significantly decreased mean burst frequency, interburst interval CV, burst duration, and percentage of spikes within bursts from 0 minutes to 4 hours. Interestingly, we found that NMDA receptors are largely responsible for neuronal network bursting, as exposure to only AP5 was enough to abolish the bursting, while blocking only AMPA receptors with NBQX only partially reduced the bursting (Fig. S3). These results agree with previous reports showing that Ketamine and Xenon which act on NMDA receptors, lead to burst silencing and reduction in vitro^40,41^. No changes in firing rate, spikes per burst, and burst duration were seen after NBQX application alone but when AP5 or NBQX+AP5 was applied, no bursts were observed therefore firing rate, spikes per burst, and burst duration were not quantifiable (Fig. S3).

### Theta-Burst Stimulation Modulated Synaptic Plasticity

To generate input specific evoked activity from electrical stimulation, theta burst stimulation (TBS) was delivered to 13-week-old organoids 4 times with 13-minute intervals between TBS (Fig. 7A) on the HD-MEA. Four to five organoids were seeded on each well and grown on the MEA until DOM 32 and 33 before stimulation (Fig. S7). The MaxWell HD-MEA has an electrode size of 8.75 x 12.50 µm², and the electrode center-to-center distance is 17.5 µm, allowing one neuron to be recorded by multiple electrodes. For input-specific synaptic plasticity, one neuron from each well was identified based on its footprint and spike-sorted neuron traces using the Axon Tracking assay in the MaxLab-live software (Fig. S5). Then, 32 electrodes focusing on a single neuron in each well were stimulated using a modified version of previously described LTP induction protocols^42–44^ (Fig. 7A). To optimize the stimulation of each neuron, electrodes along the entire neuron including the soma and axon were targeted for stimulation.

**Figure 7.**
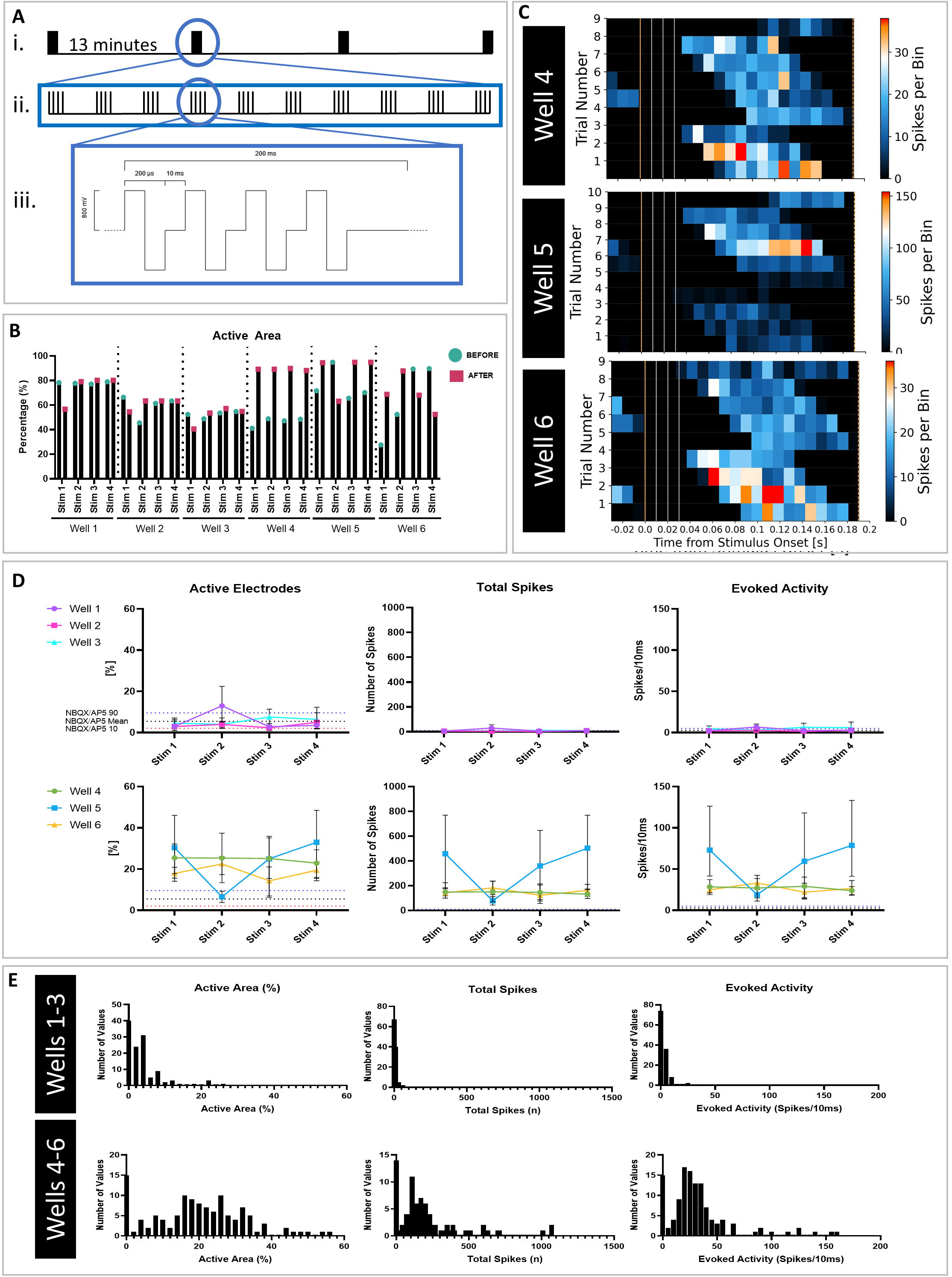
Theta-Burst Stimulation Modulated Short-Term Plasticity. A) Graphical summary of TBS protocol. i-The TBS was performed four times spaced by 13 minutes. ii-Within each TBS there are 10 trails with four spikes per trial. iii-The schematic of each trial. B) Percent active area before and after stimulation across all 6 wells. Wells 4-6 show consistent increase or decrease in active area in response to stimulation while wells 1-3 show little change. C) Representative heat map evoked activity response for wells 4-6. Bin size is equal to 10 ms. The stimulation pulses are the light grey vertical lines, and the dashed orange lines indicate the start/stop time of the analysis window for calculating evoked activity. D) Active electrodes, total spikes, and evoked activity for wells 1-3 and then 4-6. The 90^th^ percentile response of a well treated with NBQX/AP5 before and during stimulation is shown in blue overlayed on all graphs. The mean response of a well treated with NBQX/AP5 before and during stimulation is shown in black overlayed on all graphs. The 10^th^ percentile response of a well treated with NBQX/AP5 before and during stimulation is shown in red overlayed on all graphs. Responses above this NBQX/AP5 region indicate responses generated by glutamatergic receptors. E) Histograms of total evoked activity per bin (bin size of 10 ms), total spikes, and total active area. The top three graphs show data aggregated across all electrodes for all 4 TBS for wells 1-3 and the bottom three graphs show data aggregated across all electrodes for all 4 TBS for wells 4-6. Wells 1-3 show little to no response while wells 4-6 indicate evoked responses on the millisecond timescale.

To investigate short-term changes in evoked activity, total evoked activity per bin (10 ms), total spikes, and total active area were measured. Active area before and after each stimulation is shown for each well (Fig. 7B), indicating that wells 4-6 showed significant changes in active area in response to the stimulus. Representative evoked activity heatmaps for wells 4-6 indicated large responses in the milliseconds following stimulation (Fig. 7C). An interesting pattern emerged: after each TBS trial, the evoked response occurred faster until it was immediate, then returned to a longer latency, repeating this pattern across all four TBS sets for wells 4-6 but not in wells 1-3 (Fig. S9). Wells 1-3 showed no increased active electrodes, spikes, or evoked activity above threshold following TBS (Fig. 7D).

To determine an activity threshold, a separate experiment treated one well with NBQX/AP5 to block glutamatergic receptors-dependent synaptic plasticity. The 90^th^, mean, and 10^th^ percentile responses from the NBQX/AP5-treated well is shown overlayed on the plots as the dotted blue, black, and red lines, respectively (Fig. 7D). Wells 1-3 did not exceed this threshold, while wells 4-6 consistently did across all four TBS sets (Fig. 7D). Aggregated data for active area, total spikes, and evoked activity show that wells 4-6 had a bimodal distribution with increased activity, while wells 1-3 only exhibited a mode around 0 (Fig. 7E). Wells 1-3, with lower baseline activity and connectivity compared to wells 4-6, did not respond above threshold, whereas the second mode in wells 4-6 suggests short term potentiation, as stimulation lead to short term increases in activity.

Additionally, criticality and connectivity were quantified before, during, and after TBS. The BR increased during and after TBS, while the DCC median decreased, and the SCe median remained stable during TBS but decreased after TBS (Fig. S7A). These results suggest TBS caused the organoids to approach a more critical state, maintained for at least 3 hours post-TBS. Overall, functional connectivity and network dynamics remained largely unchanged over time (Fig. S7B).

Long-term effects of TBS on organoids were assessed by analyzing the number of units, total spikes, and normalized spikes across units for wells 4-6 (Fig. S7C). No consistent trends over time were observed in these metrics for wells exhibiting STP (Fig. S7C). Voltage plots before and after stimulation showed limited significant changes: well 5 had a significant decrease in maximum peak amplitude 120 minutes post-stimulation, while well 6 showed a significant increase (Fig. S7D), correlating with the number of spike-sorted units in wells 5 and 6. Interspike Interval (ISI) was calculated with a 4 Hz threshold (up to 250 ms) to account for changes in theta entrainment/phase locking. Well 4 showed mostly lower ISIs except after stimulation four (Fig. S10E). Well 5 had lower ISIs after the first stimulation throughout 180 minutes, while well 6 had higher ISIs after the first stimulation throughout 180 minutes. The coefficient of variation (CV) was used to measure ISI variability across timepoints^7^. A CV of 2.5 indicates a perfect Poisson process^7,45^, while a CV near zero indicates a perfectly periodic spike train^7,46^. Well 4 showed a significant CV decrease after stimulation four, well 5 showed a significant CV increase 180 minutes post-first stimulation, and well 6 showed a significant CV decrease 120 minutes post-first stimulation (Fig. S7F). These data indicate there are some input-specific TBS-induced changes in connected neurons over hours but not the overall network, supporting the use of this model to modulate synaptic plasticity and detect changes in synaptic plasticity in connected neurons.

## Discussion

By studying key molecular and functional changes in organoids, we aim to validate neural organoids as an *in vitro* model of learning and memory providing a human-relevant platform for translating basic science to human applications. Despite recent studies on bMPS electrophysiology, investigations into connectivity, criticality, and synaptic plasticity remain limited. Our study examined spontaneous and evoked neuronal network dynamics, functional connectivity, IEG expression, and synaptic plasticity, offering insights into learning and memory in bMPS systems. IEGs are restricted to the neurons that are engaged in learning, making their expression a prerequisite of learning capabilities. We showed the expression of *ARC*, *EGR1*, *BDNF*, *NPAS4*, *NPTX2*, and *FOS* in organoids. *ARC* and *EGR1* (also known as *Zif268, Krox-24, or NGFI-A*) are calcium-regulated IEGs essential for late LTP and long-term memory^47,48^. Additionally, CAMK2A is a key protein involved in synaptic plasticity and memory^49,50^. When CAMK2A is activated, it phosphorylates CREB allowing it to bind to the cAMP response element on the DNA^51^. *CREB*, while not an IEG, is a transcription factor vital for the expression of IEGs including *ARC* and *BDNF* ^52,53^. CAMK2 can phosphorylate SYNGAP1 mediating LTP^54–56^. Therefore, the expression of IEGs and synaptic plasticity related genes supports the potential for LTP in the neural organoids (Fig. 2C). We also showed the expression of synaptic plasticity related miRNAs. *miR-124* controls signaling molecules involved in synaptic plasticity and memory formation and *miR-132* responds to neuronal activity *in vivo* and may play a role in experience-dependent neuronal plasticity^57,58^. In contrast, *mir-134* is important for synaptic downscaling^59^ and inhibition of *mir-134* has been shown to rescue LTP^60^. Confirming this, our model showed reciprocal expression of miR-124 and miR-132 with miR-134.

Using calcium imaging, organoids exhibited spontaneous bursting starting at 4 weeks of differentiation. Calcium imaging transients showed higher frequency bursting events in week 4 to 6 organoids. At week 8, calcium transients had longer burst duration indicating sustained action potentials. These changes in calcium dynamics over time are consistent with results from dissociated rat cells cultured in 3D^61,62^. The changes in calcium transients over time could be attributed to changes in cellular populations, such as the maturation of oligodendrocytes. Oligodendrocyte populations mature after eight weeks of differentiation^11^ which is also when the largest change in calcium oscillation occurs, suggesting their contribution to these changes. The recording of Ca^2+^ transients were technically limited to 6 min, thus the absence of the oscillation at week 12 can indicate longer interburst intervals rather than absence of the activity (activity confirmed with HD-MEA recordings).

Further HD-MEA analysis showed differing network dynamics among the different age groups. When compared to 10-13 age group, the week 6-9 group had higher frequency bursting events, a larger number of nodes and edges and lower number of modularity indicating a robust and connected network of neurons; showed higher neurite outgrowth as shown in active area over time (Fig. 4C) likely contributing to the lower modularity (Fig. 5D). As organoids matured, they had fewer nodes but stronger connections, approaching a critical state, confirming system maturity. Understanding criticality in organoid models allows us to better understand the relevance and application of these models in experimental studies.

Electrical activity over time showed a high standard deviation across both groups, with a higher standard deviation in the week 10-13 group. A small subset of the 10-13 group was never active (Fig. 3B and Fig. 4C) while the majority of week 6-9 organoids remained active throughout this period (Fig. 3B and Fig. 4C). This highlights the variability of organoid’s HD-MEA electrophysiology data and implies that the sample size needs to be high enough to account for this variability.

Neural organoids responded as expected to pharmacological challenges with receptor agonists and antagonists, indicating functional synapse receptors and channels. 4-AP and bicuculline have been used in previous studies to induce chemical long-term potentiation (LTP) as they increase synaptic transmission^63^. Therefore, stimulating with 4-AP and bicuculline, then quantifying IEG expression and neuronal network activity confirmed that organoids express the molecular machinery involved in LTP. The organoids responded electrically to bicuculline, but the bulk RNA gene expression data showed no significant increase in IEG expression after bicuculline treatment. This is likely because the population of GABAergic neurons is smaller than Glutamatergic neurons in the model, making the IEG expression changes upon blockage of GABAergic neurons more difficult to detect with bulk gene expression. It’s been estimated that the ratio of GABAergic neurons to other neurons in cortical regions is 1 to 5 or 20 percent^64,65^ therefore our small population of GABAergic neurons corresponds with these estimations.

*NPAS4* gene expression is induced by calcium influx in the post-synaptic terminal after neuronal activity^66^, this correlates with our findings that 4-AP exposure caused an increase in *NPAS4* expression in both week 8 and 13 organoids. Interestingly, NPAS4 regulates the expression of multiple genes including *BDNF* and *NPTX2*^66^. While we detected an increase in *NPAS4* gene expression, we did not observe an increase in its downstream targets BDNF and NPTX2. This can be explained by the time point of sample collection (2 hours after exposure), and an increase in the downstream targets might be seen later. NBQX and AP5 disrupt excitatory glutamatergic synaptic transmission, specifically network bursting, therefore, as expected there is no change in expression of synaptic plasticity related genes after addition of these compounds.

After electrical theta burst stimulation, neuronal synaptic plasticity, connectivity and criticality were investigated. We identified candidate electrodes that showed an increase in activity immediately after stimulation and were maintained for short time periods. This approach towards teasing apart input specific STP/STD on HD-MEAs by identifying one neuron to stimulate (using 32 electrodes), allows for the determination of input specific synaptic plasticity rather than network level events that previous MEA-based studies have investigated. Longer time scale analysis indicated slight shifts in neuron level voltage, ISI, and CV values demonstrating the TBS could have input specific long-term effects on sub populations of neurons within organoids. In addition, criticality changed after TBS and drove the organoids to a more critical state. Despite this, critical dynamics after the TBS were not as pronounced as previously reported when neuronal systems are exposed to more structured stimulation in a closed-loop setup^27^. This supports the theory that neuronal criticality arises in dynamic systems when presented with structured information, maximizing information capacity and transmission through the network. Here, less structure was contained in the signal presented to the cultures compared with previous work^27^ and therefore while there was a shift towards more critical state, a greater shift might be observed if the signal contained greater complexity. These results highlight the value of considering criticality as a continuum. The ability to see nuanced changes supports using criticality as an endpoint to assess synaptic plasticity. The lower baseline activity in wells 1-3 resulted in a relative low number of active neurons after spike sorting, which can explain the absence of response after stimulation in these wells. Thus, a certain baseline activity threshold should be set up and used as acceptance/validity criteria for such experiments to increase reproducibility.

While this study demonstrates an initial attempt to examine input-specific synaptic plasticity in human neural organoids using a HD-MEA, further research using more complex models is needed. Cortical-hippocampal assembloids would be essential to study the mechanisms of learning and memory, as specific synapses in the hippocampus are pivotal for these functions^67–69^.

Combining these functional endpoints with established disease models in organoids will enhance research on disease pathophysiology, drug development, toxicant identification and various genetic, infectious, neurodevelopmental, and neurodegenerative disorders.

This study builds on the concept of OI^13^ by exploring the molecular and functional aspects of synaptic plasticity underlying learning and memory capabilities in neural organoids. An OI community is forming, which is embracing the ethical challenges of this approach^70,71^.

Future work will explore reinforcement learning in the bMPS model and in more complex bMPS models, advancing the concept of “learning-in-a-dish” towards OI.

## METHODS

### RESOURCE AVAILABILITY

#### Lead contact

Further information and requests for resources and reagents should be directed to and will be fulfilled by the lead contact, Lena Smirnova.

#### Materials availability

This study did not generate new unique reagents.

### EXPERIMENTAL MODEL AND SUBJECT DETAILS

#### Brain Microphysiological System

Female fibroblast (donor cell material: MRC-9) derived NIBSC8 (N8) iPSCs were obtained from National Institute for Biological Standards and Control, NIBSC (NIBSC), UK with a certificate of analysis identifying that they have no mycoplasma, bacteria, or viruses and have normal karyotype as identified by SNP Array. hiPSCs were cultured in mTESR-Plus medium (StemCell Technologies) at 5% O_2_, 5% CO_2_ and 37 °C. Stemness was confirmed with Oct4, Nanog, TRA-1-61, and Sox2 by immunocytochemistry and flow cytometry (Romero et al., 2023). hiPSCs cells were then differentiated in a monolayer to neuroprogenitor cells (NPCs) using a serum-free, neural induction medium (Gibco, Thermo Fisher Scientific). Nestin/Sox2-positive NPCs were then expanded and seeded in uncoated 6-well plates. These cultures were kept with neural induction medium at 37°C, 5% CO_2_, and 20% O_2_ under constant gyratory shaking (88 rpm, 19 mm orbit) to form spheres. After 48 hours, differentiation was induced by replacing the neural induction medium with brain differentiation medium: B-27™ plus kit, 1% Glutamax (Gibco, Thermo Fisher Scientific), 10 ng/ml human recombinant GDNF (GeminiBio™), 10 ng/ml human recombinant BDNF (GeminiBio™), 1% Pen/Strep/Glutamine (Gibco, Thermo Fisher Scientific). Half changes of medium were performed 3 times a week.

---

**Supplemental Figure 1.**
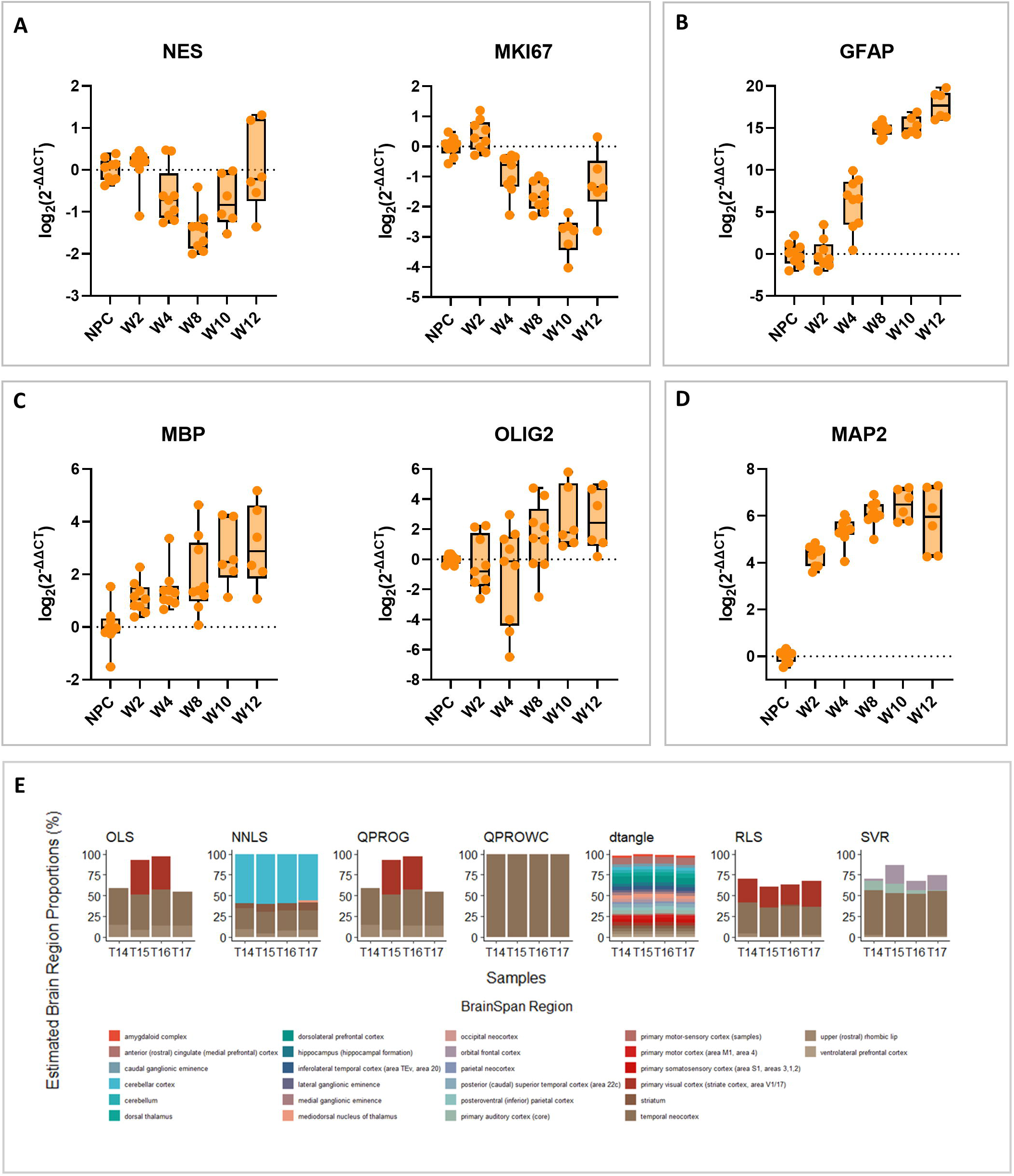
A) Expression of NPC and proliferation markers, B) astrocyte marker, C) glial markers D) neuronal dendrite marker at weeks (W) 2, 4, 8, 10 and 12 of organoid differentiation. A-D) Data is represented as log_2_ of the difference in expression, normalised to NPCs. *ACTB* was used as a reference gene. Data represents 2-3 independent experiments with 3 technical replicates each. E) Estimated brain region proportions of W8 organoids using multiple deconvolution methods (ordinary least squares, non-negative least squares regression, quadratic programming without constraints, quadratic programming non-negative and sum-to-one constraints, dtangle, robust linear regression, support vector regression) on bulk RNAseq data (4 biological replicates – T14-T17).

## Acknowledgments

We thank Dr. Paul Worley (Department of Neuroscience, Johns Hopkins University) for helpful discussions and antibody reagents. We thank George McNamara, PhD, the Ross Fluorescence Imaging Center (Johns Hopkins University), and the NIH shared instrumentation grant 1S10OD025244-01 for the use of the FV3000RS. We thank all members of the Center for Alternatives to Animal Testing for technical help and support. We gratefully acknowledge research support from the JHU SURPASS program.

D.M.A.E.D was supported by the National Institutes of Health (T32 ES007141) and International Foundation for Ethical Research Graduate Fellowship. M.S. was supported by the Deutsche Forschungsgemeinschaft (DFG, 507269789).

## Author Contributions

LS: Conceptualization, Methodology, Resources, Writing - Review & Editing, Supervision, Project Administration, and Funding acquisition. DA: Conceptualization, Methodology, Software, Validation, Formal analysis, Investigation, Writing - Original Draft. LM: Investigation and Validation. MS: Investigation and Writing - Review & Editing. JS, JL, AL, FH, TM, AM: Software, Formal Analysis, Visualization, and Writing - Review & Editing. EJ, TH, BK: Supervision, and Writing - Review & Editing. BK: Conceptualization. All authors contributed to the article and approved the submitted version.

## References

1. Smirnova, L. & Hartung, T. The Promise and Potential of Brain Organoids. Adv. Healthc. Mater. e2302745 (2024) doi:10.1002/adhm.202302745.

2. Acharya, P., Choi, N. Y., Shrestha, S., Jeong, S. & Lee, M.-Y. Brain organoids: A revolutionary tool for modeling neurological disorders and development of therapeutics. Biotechnol. Bioeng. 121, 489–506 (2024).

3. Birey, F. et al. Assembly of functionally integrated human forebrain spheroids. Nature 545, 54–59 (2017).

4. Qian, X. et al. Brain Region-specific Organoids using Mini-bioreactors for Modeling ZIKV Exposure. Cell 165, 1238–1254 (2016).

5. Quadrato, G. et al. Cell diversity and network dynamics in photosensitive human brain organoids. Nature 545, 48–53 (2017).

6. Sloan, S. A. et al. Human Astrocyte Maturation Captured in 3D Cerebral Cortical Spheroids Derived from Pluripotent Stem Cells. Neuron 95, 779–790.e6 (2017).

7. Sharf, T. et al. Functional neuronal circuitry and oscillatory dynamics in human brain organoids. Nat. Commun. 13, 4403 (2022).

8. Paşca, A. M. et al. Functional cortical neurons and astrocytes from human pluripotent stem cells in 3D culture. Nat. Methods 12, 671–678 (2015).

9. Kagan, B. J. et al. The technology, opportunities, and challenges of Synthetic Biological Intelligence. Biotechnol. Adv. 68, 108233 (2023).

10. Pamies, D. et al. A Human Brain Microphysiological System Derived from Induced Pluripotent Stem Cells to Study Neurological Diseases and Toxicity. ALTEX 34, 362–376 (2017).

11. Romero, J. C. et al. Oligodendrogenesis and myelination tracing in a CRISPR/Cas9-engineered brain microphysiological system. Front. Cell. Neurosci. 16, (2023).

12. Trujillo, C. A. et al. Complex oscillatory waves emerging from cortical organoids model early human brain network development. Cell Stem Cell 25, 558–569.e7 (2019).

13. Smirnova, L. et al. Organoid intelligence (OI): the new frontier in biocomputing and intelligence-in-a-dish. Front. Sci. 1, (2023).

14. Cai, H. et al. Brain organoid reservoir computing for artificial intelligence. Nat. Electron. 6, 1032–1039 (2023).

15. Citri, A. & Malenka, R. C. Synaptic Plasticity: Multiple Forms, Functions, and Mechanisms. Neuropsychopharmacology 33, 18–41 (2008).

16. Kotaleski, J. H. & Blackwell, K. T. Computational Neuroscience: Modeling the Systems Biology of Synaptic Plasticity. Nat. Rev. Neurosci. 11, 239–251 (2010).

17. Mateos-Aparicio, P. & Rodríguez-Moreno, A. The Impact of Studying Brain Plasticity. Front. Cell. Neurosci. 13, 66 (2019).

18. Stampanoni Bassi, M., Iezzi, E., Gilio, L., Centonze, D. & Buttari, F. Synaptic Plasticity Shapes Brain Connectivity: Implications for Network Topology. Int. J. Mol. Sci. 20, 6193 (2019).

19. Bliss, T. V. P. & Lømo, T. Long-lasting potentiation of synaptic transmission in the dentate area of the anaesthetized rabbit following stimulation of the perforant path. J. Physiol. 232, 331–356 (1973).

20. Morris, R. G., Anderson, E., Lynch, G. S. & Baudry, M. Selective impairment of learning and blockade of long-term potentiation by an N-methyl-D-aspartate receptor antagonist, AP5. Nature 319, 774–776 (1986).

21. Volianskis, A., Collingridge, G. L. & Jensen, M. S. The roles of STP and LTP in synaptic encoding. PeerJ 1, e3 (2013).

22. Volianskis, A. & Jensen, M. S. Transient and sustained types of long-term potentiation in the CA1 area of the rat hippocampus. J. Physiol. 550, 459–492 (2003).

23. France, G. et al. Differential regulation of STP, LTP and LTD by structurally diverse NMDA receptor subunit-specific positive allosteric modulators. Neuropharmacology 202, 108840 (2022).

24. Minatohara, K., Akiyoshi, M. & Okuno, H. Role of Immediate-Early Genes in Synaptic Plasticity and Neuronal Ensembles Underlying the Memory Trace. Front. Mol. Neurosci. 8, 78 (2016).

25. Chu, H.-Y. Synaptic and cellular plasticity in Parkinson’s disease. Acta Pharmacol. Sin. 41, 447–452 (2020).

26. Heiney, K. et al. Criticality, Connectivity, and Neural Disorder: A Multifaceted Approach to Neural Computation. Front. Comput. Neurosci.15, (2021).

27. Habibollahi, F., Kagan, B. J., Burkitt, A. N. & French, C. Critical dynamics arise during structured information presentation within embodied in vitro neuronal networks. Nat. Commun.14, 5287 (2023).

28. Skilling, Q. M., Ognjanovski, N., Aton, S. J. & Zochowski, M. Critical Dynamics Mediate Learning of New Distributed Memory Representations in Neuronal Networks. Entropy 21, 1043 (2019).

29. Osaki, T. et al. Complex activity and short-term plasticity of human cerebral organoids reciprocally connected with axons. Nat. Commun. 15, 2945 (2024).

30. Molen, T. van der et al. Protosequences in human cortical organoids model intrinsic states in the developing cortex. 2023.12.29.573646 Preprint at 10.1101/2023.12.29.573646 (2023).

31. Osaki, T. & Ikeuchi, Y. Advanced Complexity and Plasticity of Neural Activity in Reciprocally Connected Human Cerebral Organoids. 2021.02.16.431387 Preprint at 10.1101/2021.02.16.431387 (2021).

32. Patton, M. H. et al. Synaptic plasticity in human thalamocortical assembloids. Cell Rep. 43, (2024).

33. Zafeiriou, M.-P. et al. Developmental GABA polarity switch and neuronal plasticity in Bioengineered Neuronal Organoids. Nat. Commun. 11, 3791 (2020).

34. Developmental Transcriptomelil:: BrainSpan: Atlas of the Developing Human Brain. https://www.brainspan.org/rnaseq/search/index.html.

35. Hunt, D. L. & Castillo, P. E. Synaptic plasticity of NMDA receptors: mechanisms and functional implications. Curr. Opin. Neurobiol. 22, 496–508 (2012).

36. Bar-Shira, O., Maor, R. & Chechik, G. Gene Expression Switching of Receptor Subunits in Human Brain Development. PLOS Comput. Biol. 11, e1004559 (2015).

37. Jeyabalan, N. & Clement, J. P. SYNGAP1: Mind the Gap. Front. Cell. Neurosci. 10, 32 (2016).

38. Hu, Z. & Li, Z. miRNAs in Synapse Development and Synaptic Plasticity. Curr. Opin. Neurobiol. 45, 24–31 (2017).

39. Cocchi, L., Gollo, L. L., Zalesky, A. & Breakspear, M. Criticality in the brain: A synthesis of neurobiology, models and cognition. Prog. Neurobiol. 158, 132–152 (2017).

40. Ahtiainen, A. et al. Ketamine reduces electrophysiological network activity in cortical neuron cultures already at sub-micromolar concentrations – Impact on TrkB-ERK1/2 signaling. Neuropharmacology 229, 109481 (2023).

41. Uchida, T. et al. Xenon-induced inhibition of synchronized bursts in a rat cortical neuronal network. Neuroscience 214, 149–158 (2012).

42. Kelleher, R. J., Govindarajan, A. & Tonegawa, S. Translational Regulatory Mechanisms in Persistent Forms of Synaptic Plasticity. Neuron 44, 59–73 (2004).

43. Nguyen, P. V. & Kandel, E. R. Brief theta-burst stimulation induces a transcription-dependent late phase of LTP requiring cAMP in area CA1 of the mouse hippocampus. Learn. Mem. 4, 230–243 (1997).

44. Caneus, J. et al. A human induced pluripotent stem cell-derived cortical neuron human-on-a chip system to study Aβ42 and tau-induced pathophysiological effects on long-term potentiation. Alzheimers Dement. Transl. Res. Clin. Interv. 6, e12029 (2020).

45. Poisson Model of Spike Generation | Request PDF. https://www.researchgate.net/publication/2807507_Poisson_Model_of_Spike_Generation.

46. Maimon, G. & Assad, J. A. Beyond Poisson: Increased Spike-Time Regularity Across Primate Parietal Cortex. Neuron 62, 426–440 (2009).

47. Guzowski, J. F. et al. Inhibition of Activity-Dependent Arc Protein Expression in the Rat Hippocampus Impairs the Maintenance of Long-Term Potentiation and the Consolidation of Long-Term Memory. J. Neurosci. 20, 3993–4001 (2000).

48. Thiel, G., Mayer, S. I., Müller, I., Stefano, L. & Rössler, O. G. Egr-1—A Ca2+-regulated transcription factor. Cell Calcium 47, 397–403 (2010).

49. Takemoto-Kimura, S. et al. Calmodulin kinases: essential regulators in health and disease. J. Neurochem. 141, 808–818 (2017).

50. Zalcman, G., Federman, N. & Romano, A. CaMKII Isoforms in Learning and Memory: Localization and Function. Front. Mol. Neurosci. 11, 445 (2018).

51. Kasahara, J., Fukunaga, K. & Miyamoto, E. Activation of calcium/calmodulin-dependent protein kinase IV in long term potentiation in the rat hippocampal CA1 region. J. Biol. Chem. 276, 24044–24050 (2001).

52. Kim, J. & Kaang, B.-K. Cyclic AMP response element-binding protein (CREB) transcription factor in astrocytic synaptic communication. Front. Synaptic Neurosci. 14, 1059918 (2023).

53. Ying, S.-W. et al. Brain-derived neurotrophic factor induces long-term potentiation in intact adult hippocampus: requirement for ERK activation coupled to CREB and upregulation of Arc synthesis. J. Neurosci. Off. J. Soc. Neurosci. 22, 1532–1540 (2002).

54. Fu, Z. et al. Differential roles of Rap1 and Rap2 small GTPases in neurite retraction and synapse elimination in hippocampal spiny neurons. J. Neurochem. 100, 118–131 (2007).

55. Meili, F. et al. Multi-parametric analysis of 57 SYNGAP1 variants reveal impacts on GTPase signaling, localization, and protein stability. Am. J. Hum. Genet. 108, 148–162 (2021).

56. Zhu, J. J., Qin, Y., Zhao, M., Van Aelst, L. & Malinow, R. Ras and Rap control AMPA receptor trafficking during synaptic plasticity. Cell 110, 443–455 (2002).

57. Fischbach, S. J. & Carew, T. J. MicroRNAs in Memory Processing. Neuron 63, 714–716 (2009).

58. Nudelman, A. S. et al. Neuronal Activity Rapidly Induces Transcription of the CREB-Regulated microRNA-132, in vivo. Hippocampus 20, 492 (2010).

59. Fiore, R. et al. MiR-134-dependent regulation of Pumilio-2 is necessary for homeostatic synaptic depression. EMBO J. 33, 2231–2246 (2014).

60. Baby, N., Alagappan, N., Dheen, S. T. & Sajikumar, S. MicroRNA-134-5p inhibition rescues long-term plasticity and synaptic tagging/capture in an Aβ(1–42)-induced model of Alzheimer’s disease. Aging Cell 19, e13046 (2020).

61. Ming, Y., Hasan, M. F., Tatic-Lucic, S. & Berdichevsky, Y. Micro Three-Dimensional Neuronal Cultures Generate Developing Cortex-Like Activity Patterns. Front. Neurosci. 14, (2020).

62. Marom, A., Shor, E., Levenberg, S. & Shoham, S. Spontaneous Activity Characteristics of 3D “Optonets”. Front. Neurosci.10, (2017).

63. Jiang, Y. & VanDongen, A. M. J. Selective increase of correlated activity in Arc-positive neurons after chemically induced long-term potentiation in cultured hippocampal neurons. eNeuro 8, ENEURO.0540-20.2021 (2021).

64. Hendry, S. H., Schwark, H. D., Jones, E. G. & Yan, J. Numbers and proportions of GABA-immunoreactive neurons in different areas of monkey cerebral cortex. J. Neurosci. Off. J. Soc. Neurosci. 7, 1503–1519 (1987).

65. Sahara, S., Yanagawa, Y., O’Leary, D. D. M. & Stevens, C. F. The fraction of cortical GABAergic neurons is constant from near the start of cortical neurogenesis to adulthood. J. Neurosci. Off. J. Soc. Neurosci. 32, 4755–4761 (2012).

66. Sun, X. & Lin, Y. Npas4: Linking Neuronal Activity to Memory. Trends Neurosci. 39, 264–275 (2016).

67. Buzsáki, G. Two-stage model of memory trace formation: a role for ‘noisy’ brain states. Neuroscience 31, 551–570 (1989).

68. Fuchsberger, T. & Paulsen, O. Modulation of hippocampal plasticity in learning and memory. Curr. Opin. Neurobiol. 75, 102558 (2022).

69. Kennedy, M. B. Synaptic Signaling in Learning and Memory. Cold Spring Harb. Perspect. Biol. 8, a016824 (2016).

70. Morales Pantoja, I. E., et al. First Organoid Intelligence (OI) workshop to form an OI community. Front. Artif. Intell. 6, (2023).

71. Hartung, T. et al. The Baltimore declaration toward the exploration of organoid intelligence. Front. Sci. 1, (2023).

72. Patel, H., et al. nf-core/rnaseq: nf-core/rnaseq v3.14.0 - Hassium Honey Badger. Zenodo 10.5281/zenodo.10471647 (2024).

73. Di Tommaso, P. et al. Nextflow enables reproducible computational workflows. Nat. Biotechnol. 35, 316–319 (2017).

74. Krueger, F., et al. FelixKrueger/TrimGalore: v0.6.10 - add default decompression path. Zenodo 10.5281/zenodo.7598955 (2023).

75. Dobin, A. et al. STAR: ultrafast universal RNA-seq aligner. Bioinformatics 29, 15–21 (2013).

76. Patro, R., Duggal, G., Love, M. I., Irizarry, R. A. & Kingsford, C. Salmon: fast and bias-aware quantification of transcript expression using dual-phase inference. Nat. Methods 14, 417–419 (2017).

77. Martin, F. J. et al. Ensembl 2023. Nucleic Acids Res. 51, D933–D941 (2023).

78. Quinlan, A. R. & Hall, I. M. BEDTools: a flexible suite of utilities for comparing genomic features. Bioinformatics 26, 841–842 (2010).

79. Picard Tools - By Broad Institute. https://broadinstitute.github.io/picard/.

80. Li, H. et al. The Sequence Alignment/Map format and SAMtools. Bioinformatics 25, 2078–2079 (2009).

81. Wang, L., Wang, S. & Li, W. RSeQC: quality control of RNA-seq experiments. Bioinformatics 28, 2184–2185 (2012).

82. García-Alcalde, F. et al. Qualimap: evaluating next-generation sequencing alignment data. Bioinformatics 28, 2678–2679 (2012).

83. Sayols, S., Scherzinger, D. & Klein, H. dupRadar: a Bioconductor package for the assessment of PCR artifacts in RNA-Seq data. BMC Bioinformatics 17, 428 (2016).

84. Preseq | The Smith Lab. https://smithlabresearch.org/software/preseq/.

85. Love, M. I., Huber, W. & Anders, S. Moderated estimation of fold change and dispersion for RNA-seq data with DESeq2. Genome Biol. 15, 550 (2014).

86. Ritchie, M. E. et al. limma powers differential expression analyses for RNA-sequencing and microarray studies. Nucleic Acids Res. 43, e47 (2015).

87. Data Downloadlil:: BrainSpan: Atlas of the Developing Human Brain. https://www.brainspan.org/static/download.html.

88. sva. Bioconductor http://bioconductor.org/packages/sva/.

89. Hunt, G. J., Freytag, S., Bahlo, M. & Gagnon-Bartsch, J. A. dtangle: accurate and robust cell type deconvolution. Bioinforma. Oxf. Engl. 35, 2093–2099 (2019).

90. granulator. Bioconductor http://bioconductor.org/packages/granulator/.

91. Shannon, C. E. A mathematical theory of communication. Bell Syst. Tech. J. 27, 379–423 (1948).

92. Pachitariu, M., Sridhar, S., Pennington, J. & Stringer, C. Spike sorting with Kilosort4. Nat. Methods 21, 914–921 (2024).

93. Buccino, A. P. et al. SpikeInterface, a unified framework for spike sorting. eLife 9, e61834 (2020).

